# Acetylated microtubules are required for maintenance of the barrier between two adjacent tissues

**DOI:** 10.1101/2021.06.07.447432

**Authors:** Matthew Antel, Taylor Simao, Muhammed Burak Bener, Mayu Inaba

## Abstract

Microtubule acetylation is found in populations of stable, long-lived microtubules, occurring on the conserved lysine 40 (K40) residue of α-tubulin, catalyzed by α-tubulin acetyltransferases (αTATs). K40 acetylation has been shown to stabilize microtubules via enhancing microtubule resilience against mechanical stress. Here we show that Drosophila CG17003/leaky (Lky), an αTAT, is required for proper oogenesis. We found that loss of *lky* disrupted the cell junction between germline cyst and follicle epithelial cells, adjacent cells that form an egg chamber. This resulted in leakage of germline contents into somatic follicle cells. The follicle cells that received germline-derived nanos gene product failed to maintain their cell fate, leading to an egg chamber fusion. The same phenotype was observed upon replacement of major α-tubulin84B^K40^ with α-tubulin84B^K40A^ (non-acetylable tubulin), suggesting α-tubulin^K40^ acetylation is required for the boundary integrity of these two adjacent tissues. Taken together, this study provides the first in vivo function of tubulin acetylation in maintaining the integrity of a tissue barrier.

## Introduction

Microtubules play essential roles on various cellular processes including polarized trafficking, mitosis, migration and determining cellular rigidity [1–4]. Microtubule dynamics are regulated by post-translational modifications (PTMs) to the α- and β-tubulin subunits. One such PTM is ⍰-tubulin acetylation, which is enriched in populations of stable microtubules [5–7]. The ⍰-tubulin acetyltransferase (αTAT) enzyme responsible for acetylation of ⍰-tubulin was first identified in *Caenorhabditis elegans* as Mec-17 [8, 9] and is highly conserved across species [10, 11]. αTubulin K40 acetylation enhances resilience of microtubules against mechanical stresses [12–14]. Mutant studies in various organisms, including worms [15], zebrafish [9], mice [16, 17] and fruit flies [18], revealed that α-K40 acetylation is commonly required in mechanosensory neurons. Specific requirement of ⍰-K40-acetylation in mechanosensory neurons implies that ⍰-K40-acetylated microtubules have a qualitatively distinct role from other populations of microtubules, likely through maintaining cellular rigidity to resist external forces. While microtubules have common roles in essentially all cell types, the roles of α-K40 acetylation in non-neuronal cell types have not been well understood.

*Drosophila* oogenesis proceeds in structures called ovarioles, where egg chambers at multiple stages of development are aligned in a spatiotemporally ordered manner. The most anterior unit of the ovariole is the germarium, where germline stem cells (GSCs) continuously produce differentiating germ cells. At the tip of the germarium, GSCs divide asymmetrically to produce a cystoblast, which divides four more times and forms a 16-cell syncytium or a cyst (Figure 1A). One of the 16 germ cells within the cyst becomes the oocyte, while the other 15 cells become nurse cells, which support oocyte development [19–21]. Somatic follicle cells (FCs) are derived from stem cells residing in germarium [22–24], which give rise to precursor cells that proliferate and encapsulate the rapidly enlarging germline cyst [19, 25]. This process repeatedly occurs in adult female flies with precise spatial and temporal resolution, providing a perfect model system to study how multiple cell types interact and coordinate each other for successful organ development [19, 26].

**Figure 1.**
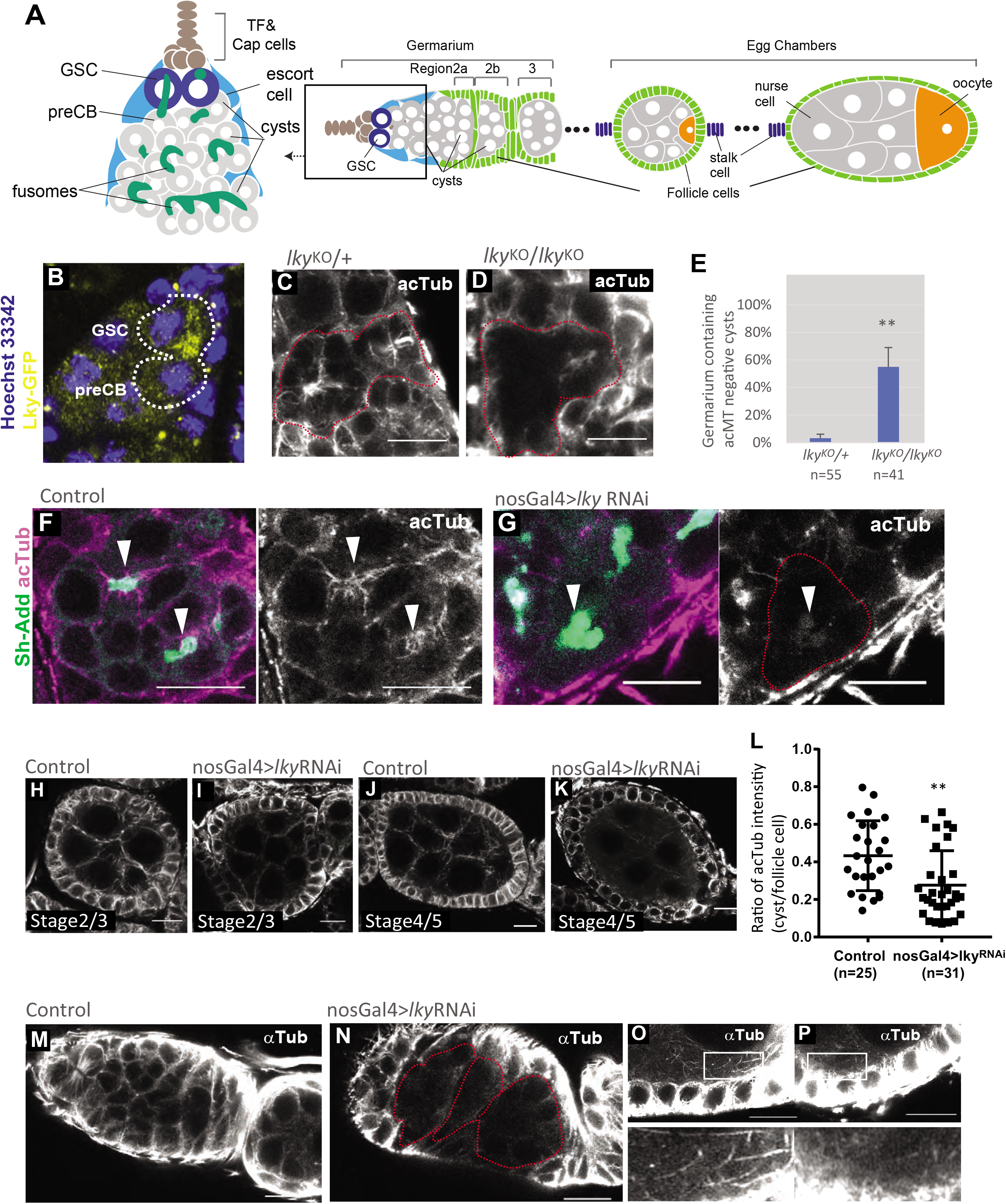
Lky is required for microtubule acetylation in germline. **A**) (Left) A schematic showing a germarium, the anterior-most unit of the ovariole which contains both germ and somatic stem cells as well as early dividing cysts. TF=terminal filament, GSC=germline stem cell, preCB=precystoblast. (Right) a schematic depicting an ovariole. Rectangle indicates the germarium shown on the left. **B**) An image of a germarium showing Lky-GFP in a GSC-preCB pair. Note: because Lky-GFP signal in the fusome was not detectable after fixation, live tissue was used for imaging. **C**) A representative image of an immunofluorescent (IF) staining of a germarium for anti-acetylated-αTubulin (acTub), showing enrichment for acTub on the fusome (*lky^KO/+^* heterozygous control). **D**) acTub is absent from the fusome in homozygous *lky*^KO^. **E**) A quantification of germline cysts lacking acTub in *lky*^KO^ ovaries. **F**) acTub staining shows colocalization with Sh-Add-GFP, a fusome marker. **G**) A germarium of germline-specific (nosGal4) knockdown of *lky* using RNAi shows no acTub staining at the fusome. **H-K**) Early stage (2-5) egg chambers show acTub in the germline in the control (**H,J**) but not in nos>*lky*RNAi ovaries (**I,K**). **L**) Quantification of acTub intensity between nos>*lky*RNAi and the control. The staining intensity of acTub was measured in the cytoplasm of germline and FCs and scored as a ratio (cyst/follicle cell). **M-N)** IF staining for αTubulin (αTub), showing cysts in nos>*lky*RNAi are negative for αTubulin. **O-P**) IF staining for αTubulin, showing that thick microtubule bundles are present cortically near the germline-FC boundary in the control (**O**), but not in nos>*lky*RNAi (**P**). Scale Bars; 10μm.

During oogenesis, microtubules play several key roles, such as oocyte determination and differentiation [27, 28]. In early stages of oogenesis, microtubules associate with a specialized germ cell-specific endoplasmic reticulum-like organelle called the fusome, where the cyst forms major microtubule organizing center (MTOC) [29]. The fusome branches through dividing germ cells within the cyst [30, 31]. The microtubules emanating from this MTOC are polarized in a manner that directs trafficking [32–34]. In later stages of oogenesis, microtubules are involved in the transport of materials into maturing oocyte from nurse cells [35–42]. Microtubules are also playing general roles in ovary, such as establishing and maintaining the cell-cell junctions, adherens junctions and tight junctions, which are essential for maintaining barrier integrity of epithelia (reviewed in [43]). Microtubules also contribute to determining cellular rigidity cooperating with actomyosin cytoskeleton [44, 45].

In the ovary, acetylated α-tubulin has been reported to be enriched on fusomes in the dividing germ cells of early oogenesis [46]. The role of α-K40 acetylation in oocyte development has been completely unknown. In this study, we demonstrate that α-K40 acetylation of microtubules is required for maintenance of the tissue boundary between germline cysts and surrounding FC epithelia. Loss of α-K40 acetylation shows breakage of cell boundary between the germline and surrounding FCs. This is accompanied by the diffusion of germline contents into FCs. We provide evidence that acetylated microtubules are required for maintaining the barrier of two adjacent tissues, preventing the collapse/breakage of the cell-cell boundary.

## Results

### Lky is required for microtubule acetylation in germline

Lky is predicted to be an αTAT based on sequence homology. We found that Lky-GFP expressed under the control of the nosGal4 driver (nosGal4>UAS-Lky-GFP) localizes to the fusome, which is enriched for acetylated microtubules [46](Figure 1B). Anti-Lky antibody staining also revealed localization of endogenous Lky in the fusome (Figure S1B), and Lky localization at the boundary of cyst and somatic cell layers (Figure S1C-D).

To examine whether Lky is responsible for microtubule acetylation at the fusome, we generated a lky knock-out fly (*lky^KO^*) in which the entire lky coding region is deleted (see Methods, Figure S1A). Anti-Lky staining was completely absent in *lky^KO^* ovaries, confirming a loss of Lky in the KO (Figure S1C-D).

Consistent with the previous report [46], we observed acetylated microtubules enriched in the fusome in control germaria (Figure 1C). In contrast, both in *lky^KO^* and *lky* RNAi, levels of acetylated α-tubulin (acTub) were reduced at the fusome (Figure 1D–G), as well as throughout the cells in the cyst (Figure 1H–L). Absence of acTub was also observed in more mature egg chambers, where the fusome had disintegrated (Figure 1H–L). These results suggest that Lky is generally required for microtubule acetylation in the *Drosophila* female germline.

As *lky^KO^* ovaries maintain acTub in somatic cells (Figure S1F), we examined if the other known αTAT, dTAT (CG3967) [18] is responsible for microtubule acetylation in somatic cells. Germ cells in *dTAT^KO^* germaria maintained acetylated fusome microtubules (Figure S1G), indicating that the germline depends on Lky but not on dTAT for α-K40 acetylation. In contrast, we observed loss of acTub in somatic cells of *dTAT^KO^* ovaries (Figure S1G). Consistently, the dTAT and lky double mutant (*Double KO*, Figure S1H) showed the anterior region of the germarium completely absent for acTub. These observations suggest a role for dTAT in acetylating the microtubules of early somatic cells and Lky in acetylating microtubules of the germline. We still observed acetylated microtubules in FCs in the posterior of germaria and later staged egg chambers and a low amount in germline in *Double KO* ovaries (Figure S1I-J), possibly indicating the existence of a third αTAT responsible for α-K40 acetylation in late-stage FCs.

We next examined how the loss of α-K40 acetylation impacts entire cellular microtubules. Anti-αTubulin antibody staining revealed a reduction of microtubules in germline cysts expressing *lky* RNAi (Figure 1M–N). Closer observation revealed an absence of long, thick microtubule bundles at the FC-germ cell boundary where other microtubules were still present (Figure 1O–P). These data suggest that Lky-specifically regulates stability of certain population of microtubules via α-K40 acetylation.

### Germline contents leak into follicle cells in *lky^KO^* ovaries

What is the consequence of the loss of acetylated microtubules in the developing germline? Although ovarioles from young *lky^KO^* females appeared mostly normal, starting ~7 days of age, we found that egg chambers often contained FCs that become positive for Vasa, a germline-specific protein, as well as the FC marker Traffic Jam (TJ) (Figure 2A–D). Vasa positive FCs were observed in any stage, but most frequently seen in germaria to stage2-3 chambers (Figure 2A, B, E). The frequency of ovarioles containing FCs positive for Vasa progressively increased with age (Figure 2F). Vasa leakage was significantly rescued by germline-specific (but not somatic cell-specific) expression of *lky* (Figure 2G), suggesting that loss of lky function in the germline caused this phenotype.

**Figure 2.**
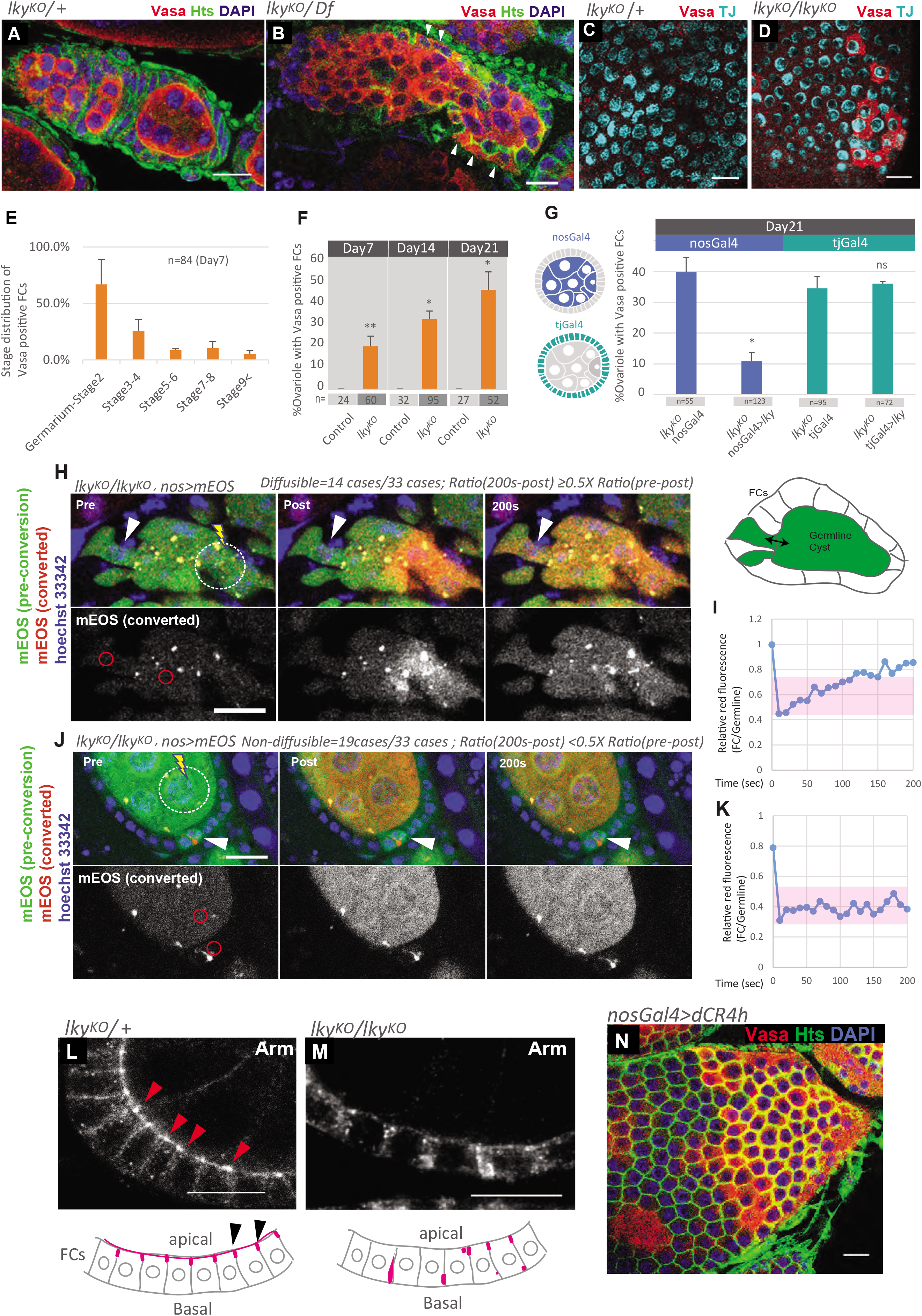
Germline contents leak into follicle cells in lkyKO ovaries. **A, B**) IF staining for Vasa and Hts (marking FC cell cortex) with DAPI (nuclei) showing abnormal Vasa staining in FC layer (white arrowhead) in *lky*^KO^ germarium. **C, D**) IF staining for Vasa and TJ showing Vasa is present in the cytoplasm of TJ-positive FCs in *lky*^KO^, looking at the follicle cells on the egg chamber surface. **E**) Quantification of the distribution of egg chamber stages containing Vasa positive FCs. Note: some ovarioles scored contained more than one egg chamber with Vasa positive FCs. **F**) A quantification of ovarioles showing ectopic Vasa localization in FCs, comparing control versus *lky*^KO^ for 7, 14, and 21 days. Percentages of ovarioles containing any follicle cells which stain positive for Vasa (Vasa in FC cytoplasm) were quantified. **G)** Germline-specific expression of *lky* in germline (under the *nosGal4* driver) reduced the frequency of ovarioles containing Vasa-positive FCs, but somatic cell-specific expression of *lky* (under the *tjGal4* driver) had no effect. **H-K**) Representative photoconversion experiments of mEOS expressed in the germline of *lky^KO^* background. White arrowhead indicates a FC positive for mEOS. mEOS was photoconverted from green to red in the germline (white circle). Red signal was monitored at red encircled areas. Equilibration of green and red signal was observed after 200 seconds. The indicated areas (red circles in **H, J,** respectively) were used for measuring the ratio of red fluorescent signal in follicle cells (FCs) to germline. Right panel shows schematic of anatomy, Pre=pre-photobleaching, Post=post-photobleaching. Change of relative intensity of red fluorescence after photoconversion is plotted in **I** and **K**. Greater than 50% of recovery (beyond the pink zone in graph **I**) was judged as “diffusible” (**H** and **I**). Whereas less than 50% of recovery (within pink zone in graph **K**) was judged as “non-diffusible” (**J** and **K**). **L-M**) IF staining for Vasa and Armadillo (Arm), a marker for adherens junctions, in the control (**L**) versus *lky^KO^* (**M**). **N**) Staining for Vasa, Hts (marking FC cell cortex), and DAPI (nuclei) in ovaries expressing dominant-negative DE-Cadherin (dCR4h) in germline (under the nosGal4 driver). Scale Bars: 10 μm.

The presence of germline contents in FCs suggested three possibilities; First, loss of microtubule acetylation led to disruption of the physical barrier between germline and FCs, resulting in “leakage” of germline cytoplasmic contents into neighboring FCs. Second, loss of acetylated microtubules in the germline leads to aberrant intercellular signaling from germline to FCs, resulting in expression of Vasa by FCs. Third, loss of acetylated microtubules may deform the germline cyst such that it extends into FCs. To distinguish these possibilities, we conducted a photobleaching assay. When mEOS was expressed using a germline-specific promoter in the *lky^KO^* background, we often saw that some FCs also expressed mEOS. To test whether this is due to the leakage of cytoplasmic content or not, we photoconverted mEOS to red specifically in germ cells. After photoconversion, we observed that converted red signal from germ cells and unconverted green signal in FC quickly mixed, reaching equilibrium within a few minutes (Figure 2H, I, movie S1) in 14/33 cases, suggesting that germline-derived mEOS can diffuse into FCs. In 19/33 cases, we observed no equilibrium after photoconversion (Figure 2J, K, movie S2), indicating that leakage in these FCs may eventually stop, and/or that FCs which received germline contents may proliferate and generate progeny that have intact cellular barriers, while maintaining the inherited “germ cell-like” cell identity. Nevertheless, we could not deny the possibility of the involvement of altered signal transduction. As a control, we expressed GFP in wildtype flies in both germ and FCs and photobleached the germline cyst, observing no recovery (Figure S2A, movie S3). This indicates that the contents of the germline and FCs do not usually diffuse into one another.

What is the cause of leakage between germline cells and FCs? During egg chamber formation, a monolayer of FCs surrounds germ cells. FCs polarize with respect to the germ cells, and the ‘apical’ side faces to the germ cells, where adherens junctions (AJs) are formed at the FC-FC boundary (Figure 2L arrowheads) [47]. In early stages of oogenesis (<Stage5), germ cells and FCs closely adhere to each other. The identity of junctions formed at the germline-FC boundary is less clear at the ultrastructural level [48]. However, several studies suggest that germline-FC attachment also uses cadherin-based adhesion [49, 50], indicating that an AJ-like structure may be also adhering germline and FC. Armadillo (Arm, a component of AJ) also localizes along the germline-FC boundary in the control (Figure 2L, bottom pink lines), but the localization was severely disrupted in *lky^KO^* (Figure 2M). We also observed AJ is often mislocalized in FCs to the lateral/basal side (Figure 2M), suggesting that apical-basal polarity of FCs is compromised. These results led us to hypothesize that disrupted cell adhesion between germline and FCs may cause germ cell cytoplasm to leak into FCs.

Indeed, expression of dominant-negative E-cadherin in germ cells (nos>dCR4h) was sufficient to cause the leakage of Vasa into FCs (Figure 2N), suggesting that disturbance of adhesion between germline and FC epithelia causes cytoplasmic leak between germ cell and FCs. It should be noted that we also observed an intercalation of the germ cell cytoplasm into the FC layer in *lky*^KO^ ovaries, likely due to compromised junction between germ cells and FCs (Figure S2B).

In summary, we concluded that Lky function is required for preventing the localization of germline contents in FCs, likely by preventing the leakage of cytoplasmic contents between germ cells and FCs.

### Microtubule defects disrupt the barrier between germline and follicle cells

To test if the observed leakage of germline contents was due to a reduction of acetylated microtubules, we incubated egg chambers in the presence of the microtubule-depolymerizing drug colcemid. Indeed, treatment of wildtype ovaries with colcemid for 30 minutes was sufficient to cause the leakage of germline contents into FCs, visualized by Vasa staining (Fig. 3A–C). These results show that depolymerizing microtubules is sufficient to disrupt the barrier between the germline and follicle cells.

**Figure 3.**
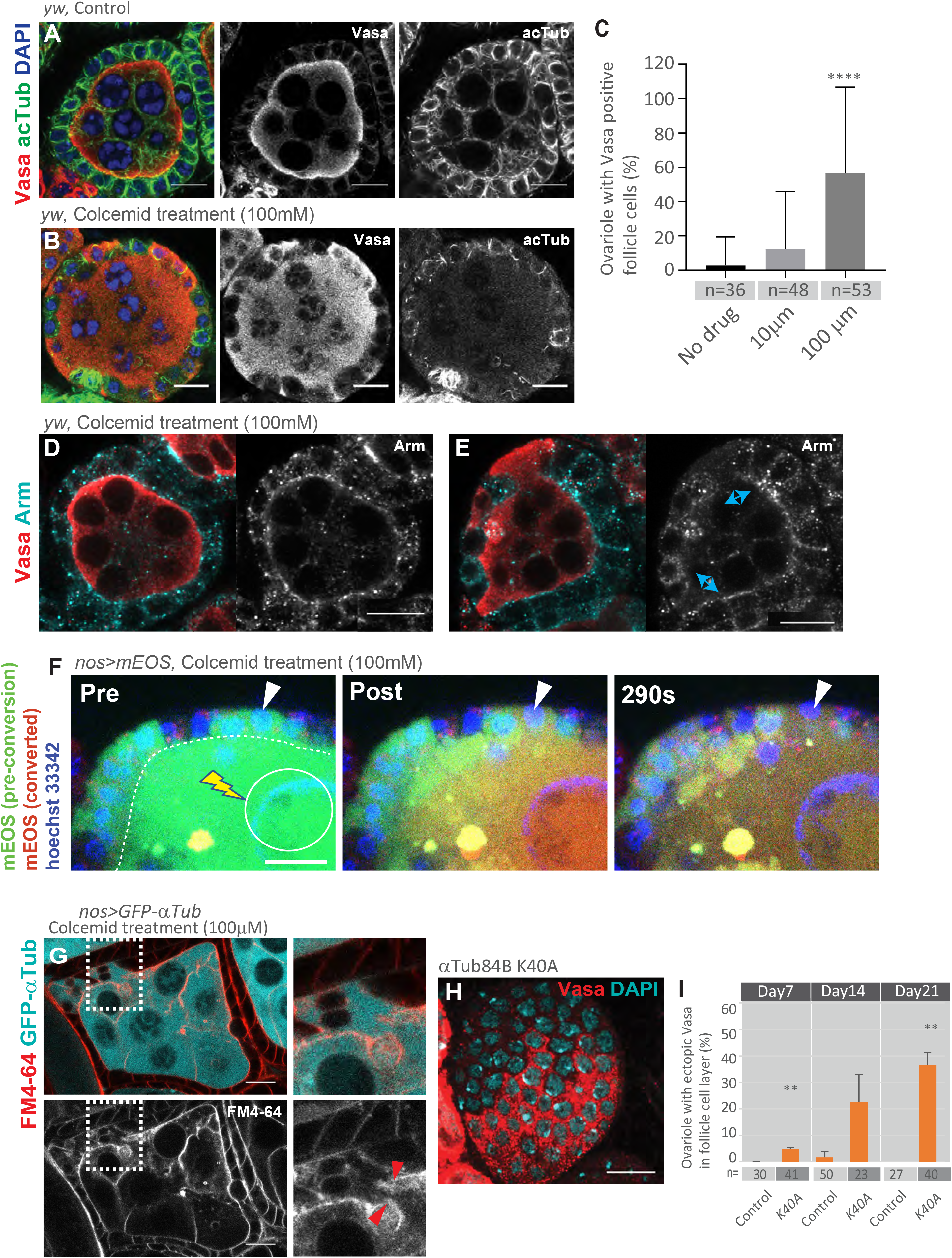
Microtubule defects disrupt the barrier between germline and follicle cells. **A, B**) Representative images of Vasa and acTub staining in stage3-4 egg chambers treated without (**A**) or with (**B**) 100 μM colcemid for 30 min to depolymerize microtubules. **C**) Quantification of the frequency of ovarioles containing Vasa-positive FCs after the indicated concentration of colcemid treatment for 30 min. **D, E**) IF staining for Vasa and Arm in control ovaries versus those treated with colcemid reveals disruption of Arm at the boundary of germ cells and Vasa-positive FCs (blue arrows). **F**) A representative photoconversion experiment after colcemid treatment. mEOS was expressed in the germline using *nosGal4* and was observed in the FCs post colcemid treatment. mEOS was photoconverted (red signal) in the germline, and red signal was observed diffusing into FCs (white arrowhead). **G**) A representative image of a live egg chamber (stage3-4) visualizing its plasma membrane by FM4-64 dye after colcemid treatment. Germline expressed GFP-□Tub shows leakage into FCs through ruptured plasma membrane. Right panels show magnified portion of missing membrane (squared in left panels). **H**) αTub84B^K40A^ knock-in ovaries show Vasa-positive follicle cells. **I**) Quantification of ovarioles containing Vasa-positive follicle cells in αTub84B^K40A^ at days 7, 14, and 21 post-eclosion.

Staining colcemid-treated ovaries for Arm revealed that leakage of Vasa into somatic cells was associated with the absence of Arm staining on the germ cell-FC boundary (Figure 3D, E). Moreover, visualizing plasma membrane by FM4-64 membrane dye revealed the absence of plasma membrane on the germ cell-FC boundary during leakage (Figure 3G). In addition to leakage of cytoplasmic contents, we also often observed an intercalation of the germ cell cytoplasm into the FC layer after colcemid treatment as seen in *lky^KO^* ovaries (Figure S3A), suggesting the junction between germ cells and FC boundary is weakened upon loss of microtubules.

Similar to the case of *lky*^KO^ egg chambers, we detected exchange of cytoplasmic contents between germ cells and FCs after colcemid treatment. When mEOS was expressed using the germline driver *nosGal4*, photoconverted red signal from the germline was observed in the FCs, suggesting diffusion of mEOS from the germline into FCs (Figure 3F, movie S4), suggesting that microtubules are required to maintain the barrier between germline cysts and FC epithelia.

Vasa+ FCs were also observed in a homozygous αTub^K40A^ knock-in line, where αTub84B gene, the most abundant αTub, is replaced with αTub84B^K40A^ in which the K40 residue is mutated to alanine (A) such that the tubulin cannot be acetylated (Figure 3H, I) [51]. This indicated that the loss of α-tubulin acetylation is the cause of the leakage of germline components into somatic FCs.

These results suggest that the disturbance of microtubules results in disruption of the junction and plasma membrane between germline cysts and FC epithelia, leading to a leakage of cytoplasmic contents.

### ⍰-K40 acetylation by Lky is required for proper egg chamber development

Oogenesis proceeds in units called ovarioles, where egg chambers consisting of a germline cyst surrounded by a follicular epithelium are aligned in a spatiotemporal order. Each egg chamber is separated from the next by a string of somatic stalk cells (Figure 1A) [52–56].

Although *lky*^KO^ ovariole structure appeared normal in young animals (0-7 days after eclosion), we often observed fusion of egg chambers in older *lky*^KO^ ovaries. The phenotype worsened as the animal aged (Figure 4A–B, G). At day 14, germline cysts started showing incomplete separation of egg chambers. In germarium through region 3 cysts, we frequently observed fused egg chambers, where 2 or more cysts were encapsulated by a single FC layer. For example, in *lky^KO^* egg chambers, a single FC layer was found encapsulating 30 nurse cells and 2 oocytes (Figure 4D,F) while wildtype egg chambers contain only 15 nurse cells and one oocyte (Figure 4C–F).

**Figure 4.**
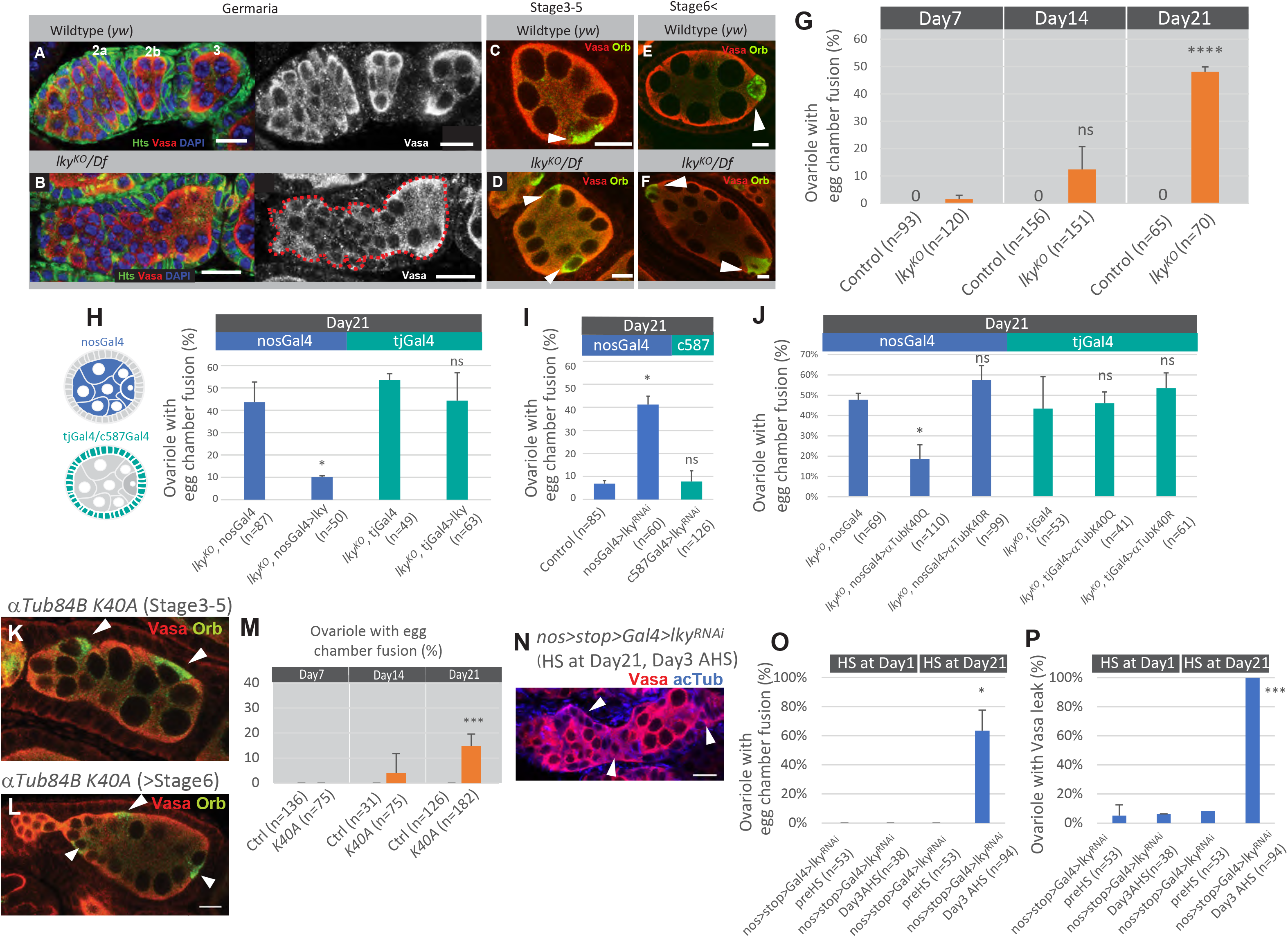
⍰-K40 acetylation by Lky is required for proper egg chamber development. **A-B)** A comparison of wildtype (**A**) and *lky*^KO^ (**B**) ovarioles, showing the egg chamber fusion phenotype of *lky*^KO^. **C-F**) Images showing oocytes (white arrowheads) in egg chambers of wildtype (**C, E**) and *lky*^KO^ (**D, F**) ovarioles. **G**) Quantification of the frequency of ovarioles with egg chamber fusion, scored at 7, 14, and 21 days. **H**) Quantification of the frequency of ovarioles with egg chamber fusion after germline-specific (nosGal4) and somatic cell-specific (tjGal4) expression of *lky*. **I**) Quantification of ovarioles with egg chamber fusion in ovaries expressing *lky* RNAi under the germline driver nosGal4 or somatic cell driver c587Gal4. **J**) Quantification of ovarioles with egg chamber fusion after germline (nosGal4) driven or somatic cell (tjGal4) driven expression of αTubulin^K40Q^ and αTubulin^K40R^ transgenes. **K-L**) αTub84B^K40A^ knock-in ovarioles showing egg chamber fusion at the indicated stage. **M**) Quantification of ovarioles with egg chamber fusion in αTub84B^K40A^ knock-in ovaries. **N)** A representative image of ovary of *lky* RNAi induced by heat shock (HS) after aging animals for 21 days. White arrowheads indicate severe “leak” of Vasa into somatic cells. **O)** Quantification of ovarioles with egg chamber fusion after heat shock at the indicated day. **P)** Quantification of ovarioles with Vasa leakage into FCs after heat shock at the indicated day. Scale Bars: 10 μm.

Expression of Lky using a germline specific driver (nosGal4), but not a somatic cell driver (tjGal4), rescued the egg chamber fusion phenotype of *lky^KO^* ovaries, and the knockdown of *lky* in the germline, but not in the somatic cells, showed the egg chamber fusion phenotype (Figure 4H, I). These data indicate that *lky* is required in the germline for normal egg chamber development.

To test if the egg chamber fusion phenotype is caused by defective K40 acetylation, we attempted to rescue the aberrant egg chambers with an α-tubulin K40Q transgene (UASp-αTub^K40Q^) in which lysine (K) 40 residue of α-tubulin84B (the most abundant α-tubulin isoform in Drosophila [57]) is mutated to glutamine (Q). α-tubulin K40Q mutation has been shown to mimic acetylated α-tubulin[9, 58, 59]. Using *nosGal4* and *tjGal4*, we drove cell type-specific expression of αTub^K40Q^ in the *lky^KO^* background. We found that germline-specific (but not somatic cell-specific) expression of αTub^K40Q^ rescued the egg chamber fusion in *lky^KO^* ovaries (Figure 4J). In contrast, a non-acetylable α-tubulin84B K40R (αTub^K40R^), which has lysine (K) 40 mutated to arginine (R), was not able to rescue the aberrant egg chamber defect (Figure 4J). Moreover, mutant flies in which the lysine (K) 40 residue of the endogenous α-tubulin84B gene is replaced to alanine (A) (αTub^K40A^) [51] exhibited a similar phenotype to *lky^KO^* flies, often showing fused egg chambers which worsened with age (Figure 4K–M).

These data indicate that the egg chamber defect observed in *lky^KO^* is due to an impairment of ⍰-K40 acetylation, and that ⍰-K40 acetylation is required for proper egg chamber development. Interestingly, when we induced *lky* RNAi (via heat shock) in aged animals, both the egg-chamber fusion and Vasa positive FCs occurred within 3 days (Figure 4N–P), indicating that age-dependent physiological changes may play a role in the establishment of this phenotype.

### Leakage of nanos (nos) gene products into FCs causes egg chamber fusion

A previous study has demonstrated that misexpression of the germline component *nanos (nos)* in FCs is sufficient to cause the phenotype of egg chamber fusion [60]. In their study, nos overexpression in FCs was sufficient to cause egg chamber fusion, and introducing a hypomorphic allele of *nos* (*nos^L7^/nos^BN^*). into the *L(3)mbt* mutant background suppressed the egg chamber defect, suggesting a direct role for somatic *nos* expression in causing egg chamber fusion.

In *lky*^KO^ ovaries, FCs positive for germline-expressed mEOS contained nos mRNA (Figure 5A, B). Therefore, we speculated that nos mRNA leak into FC may cause the phenotype of egg chamber fusion. To test whether *nos* in FCs is responsible for the egg chamber fusion in *lky*^KO^ ovaries, we analyzed ovaries from flies homozygous for both *lky*^KO^ and hypomorphic alleles of *nos* (*nos^L7^/nos^BN^*). Strikingly, we observed a significant decrease in the frequency of egg chamber fusion in these flies (Figure 5C), and overexpression of *nos* in FCs using the *c587Gal4* driver caused egg chamber fusion (Figure 5D), consistent with the previous report [60]. Together, these data suggest that *nos* gene product leaks from the germline into somatic FCs in *lky*^KO^ ovaries, resulting in egg chamber fusion.

**Figure 5.**
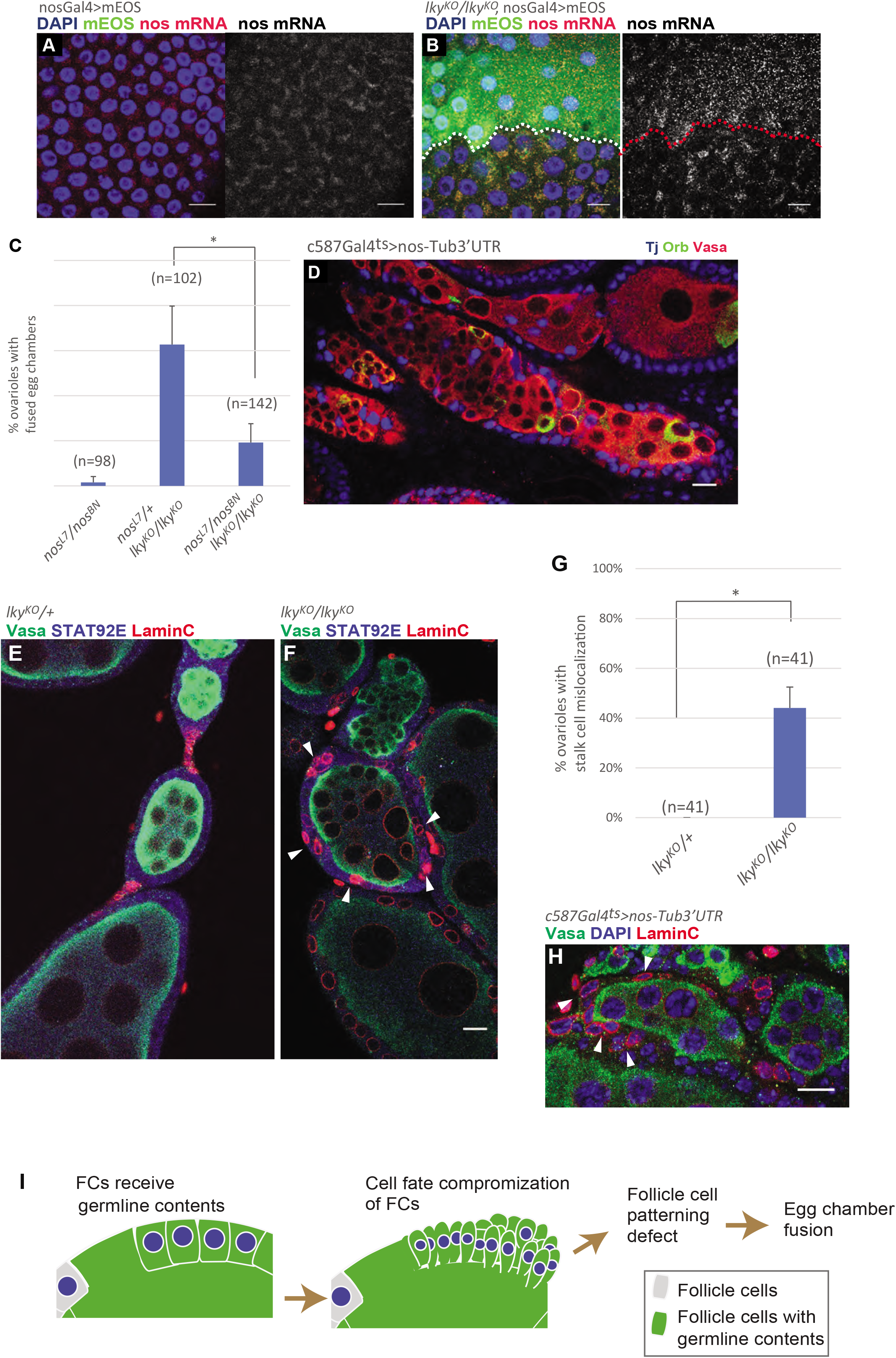
Leakage of nanos (nos) gene products into FCs causes egg chamber fusion. **A-B)** mEOS was expressed in the germline of control **(A)** and *lky^KO^* **(B)** ovaries and fluorescent in situ hybridization was performed probing for *nanos* (*nos*) mRNA. In **(B)**, mEOS from the germline is present in the FCs, and mEOS-positive follicle cells contain *nos* mRNA. **C)** A chart showing that homozygous expression of hypomorphic alleles of nos (nos^L7^ and nos^BN^) in the *lky^KO^* background rescues the egg chamber fusion defect. **D)** Expression of nos (with tubulin 3’UTR to prevent degradation) in somatic cells using c587-Gal4; Tub-Gal80^ts^ causes egg chamber fusion. **E-F)** IF staining for Vasa, Stat92e, and LaminC, showing stalk cells marked by LaminC and Stat92e expression in control **(E)** versus a fused egg chamber in *lky*^KO^ **(F)**. **G)** Quantification of the frequency of ovarioles showing stalk cell mislocalization. **H**) Representative image of LaminC positive cells mislocalizing when nos is expressed in the somatic cells. **I)** A model for germline leakage causing egg chamber fusion in *lky*^KO^ ovaries. Contents from the germline leak into FCs and reprogram them to a different cellular identity, causing a defect in their specification and patterning, leading to egg chamber fusion. Scale Bars; 10μm.

It has been reported that the egg chamber fusion is caused by a defect in the specification of stalk cells [61], which are the somatic cells connecting separate egg chambers. Notch and JAK/STAT pathways are sequentially required for FCs to differentiate into stalk cells [55, 56, 61]. In young ovaries (days 0-7) before the fusion phenotype starts appearing, we observed normal Stat92e staining in *lky*^KO^ ovarioles (Figure S5A-B), indicating that the these signaling pathways are intact. However, in fused egg chambers in aged ovaries (day 21), Lamin-C positive stalk cells were often mislocalized (Figure 5E–G), suggesting the possibility that transfer of *nos* gene product may affect stalk cell patterning, resulting in the egg chamber fusion phenotype. Consistently, overexpression of *nos* in FCs caused stalk cell mislocalization (Figure 5H).

Based on these data, we propose that loss of acTub in germline causes a leak of germline contents into FCs. FCs that receive *nos* gene product are compromised in their cell fate, which causes aberrant egg chamber development (Figure 5H).

## Discussion

The mechanism of how an epithelial layer adheres to other tissue and maintains the boundary integrity is not well understood. Here we show that α-tubulin K40 acetylation is required in the Drosophila female germline for maintaining the boundary between the germline and surrounding follicular epithelium. Without microtubule acetylation, adherens junctions become collapsed or delocalized, leading to a rupture of the plasma membrane and “leak” of germline contents into follicular epithelial cells. The leakage of germline components, especially *nanos*, into the FCs, compromises their cellular identity and leads to an egg chamber fusion.

α-Tubulin K40 acetylation has been studied extensively, yet the role of K40 acetylation in a cell has still been unclear. Since the site of acetylation, lysine (K)40 of ⍰-tubulin (⍰-K40), is located in the lumen of the microtubules [9, 62], the acetylated ⍰-K40 does not seem to be specifically recognized by other microtubule associating proteins. Therefore, after its discovery in the 1980s [5, 7, 63], it has long been unclear if the acetylated ⍰-K40 is just a passive mark of stable microtubules or if acetylation actively modulates the function of microtubules. Only recently, in vitro studies demonstrated that acetylation indeed adds specific function to microtubules, such that enhances resilience against mechanical stresses [12–14].

Our work, for the first time, demonstrates the role of acetylated microtubules in maintaining tissue boundary integrity. A recent study demonstrated that germline cysts actively generate contractile actomyosin forces, which directionally opposes the constriction force from FC layer. A proper balance of forces between germline cyst and FC constriction is required for successful encapsulation [49]. Our work suggests the possibility that the acetylated microtubules may help resolve the unbalanced forces during this process. Acetylated microtubules may act as a “cushion” when cells are under strong force to prevent the breakage of cell-cell boundary. It is tempting to test whether the role of *α*-K40 acetylation in tissue boundary maintenance seen in *Drosophila* oogenesis is common in other systems.

In many organisms, the αTAT loss-of-function phenotype is only found in mechanosensory neurons [8, 16, 18, 58]. There is a possibility that multiple αTATs function redundantly in other tissues, making it difficult to detect the phenotype. Moreover, there is also the possibility of other mechanisms at play which modulate tissue boundaries and barrier function, which could obscure any phenotype of acTub loss in other tissues. Supporting this idea, reported neuronal defects caused by loss of αTAT often worsen as the animal ages[64]. Similarly, the egg chamber fusion defect observed in *lky*^KO^ flies becomes more prevalent with age. Therefore, physiological changes in aged animals must play a role in the penetrance of this phenotype in flies defective for α-tubulin acetylation. Consistent with this idea, we observed the phenotype in aged animals within 3 days after inducing *lky* RNAi. In contrast, when *lky* RNAi started at day 0, egg chamber fusion phenotype appeared 14 days or later Identifying the “aging” factor that affects the penetrance of this phenotype would be a fascinating future study.

In summary, our data demonstrate that *α*-K40 acetylated microtubules are required for the maintenance of tissue boundary integrity. We also suggest the possibility that the disruption of the tissue boundary may increase the risk of changing cell identity due to transfer of contents between other cell types.

## Methods

### Fly husbandry and strains

All fly stocks were raised on standard Bloomington medium at 25°C. RNAi flies were placed at 29°C after eclosion for desired days before dissection. Flies used for aging were placed into new food tubes every 3-5 days. The following fly stocks were used: α*Tub84B^K40A^* (FBal0345033, [51] αTub84B knock-in, gift from J. Wildonger), *dTAT^KO^* (Fbal0345149, [18], gift from J. Wildonger), *nosGal4* (gift from YM. Yamashita)*,), sh-adducin-Venus* (gift from YM. Yamashita)*,ActGFP* (gift from YM. Yamashita)*, tjGal4* (gift from YM. Yamashita), *c587-Gal4* (gift from YM. Yamashita), *nos^L7^* (gift from R. Lehmann), *nos^BN^* (gift from R. Lehmann), *UAS-dCR4h* (gift from YM. Yamashita, [65]), *UAS-nosTub3’UTR* (gift from YM. Yamashita), and *hs-FLP; nos-FRT-stop-FRT-Gal4* (gift from YM. Yamashita, [66]. The following stocks were obtained from the Bloomington stock center Df(1)BSC586 (25420).

### Generation of transgenic and knock-out lines

#### LkyKO

*LkyKO* knockout flies were generated using the CRISPR/Cas9 system to delete the entire CG17003 coding region, as described previously [67]. Homology arms were generated using the following primer sets:

EcoRI 5’ arm Forward: CTGAATTCATATACCCAGGATGCTAGAGGGTTA
NotI 5’ arm Reverse: CTGCGGCCGCGCAGTCGCACTTAGTTCGTTTTCAT
BglII 3’ arm Forward: CTAGATCTCTGTAACTTGAGGTCTCGAACTATT
PstI 3’ arm Reverse: CTCTGCAGGGAAACTTACAAAAATTTAAGAGGC

Homology arms were then inserted into the pHD-DsRed-attP donor vector ([67], gift from M. Buszczak) by respective restriction enzyme digest and ligation.

gRNA sequences were as follows:

BbsI 5’ gRNA sequence: CTTCGAACTAAGTGCGACTGCATA BbsI
3’ gRNA sequence: CTTCGAGACCTCAAGTTACAGCCC

gRNAs were inserted into the pU6-BbsI-gRNA vector ([67], gift from M. Buszczak) by BbsI digestion and ligation. *Lky^KO^* CRISPR constructs were injected into Cas9-expressing embryos by BestGene, Inc. Recombinant flies were crossed with *w1118* and selected by expression of *3xP3-DsRed* (visualized as red eye fluorescence) and validated by genomic PCR.

Primer sequences for validation are as follows:

5’ arm:

5’-TGTGATTTGCGAATGGGATG-3’
5’-CCACCACCTGTTCCTGTA-3’

3’ arm:

5’-CTTCGAGCCGATTGTTTAG-3’
5’-ACACCTTGGAGCCGTACTGGAACT-3’

### *Lky* RNAi

For construction of Lky shRNA transgene, the following oligonucleotides were used:

<u>Line 1:</u>

5’-
ctagcagtGCGAAATCCTAAACATCATGGtagttatattcaagcataCCATGATGTTTAGGATTTCGCgcg
-3’;
5’-
aattcgcGCGAAATCCTAAACATCATGGtatgcttgaatataactaCCATGATGTTTAGGATTTCGCactg -
3’
<u>Line 2:</u>

5’-
ctagcagtGGTAGAGCCCGAGAATTATATtagttatattcaagcataATATAATTCTCGGGCTCTACCgcg
-3’;
5’-
aattcgcGGTAGAGCCCGAGAATTATATtatgcttgaatataactaATATAATTCTCGGGCTCTACCactg -
3’

Annealed oligonucleotides were subcloned into NheI and EcoRI sites of pWALIUM20 vector (Gift from Yukiko Yamashita). Transgenic flies were generated using strain attP2 by PhiC31 integrase-mediated transgenesis (BestGene).

#### UAST-αTubK40Q

For construction of UAST-GFP-*α*Tub84B K40Q, site directed mutagenesis was performed using primers:

5’-GCGGAGGTGATGACTCGTTCAACACCTTC-3’,

5’-CCACGGT<u>TTG</u>GTCAGACGGCATCTGGCCAT-3’ (K40 site is underlined) from UASP-GFP-*α*Tub84B plasmid (kind gift from Spradling A). Resultant plasmid was used to amplify *α*Tub84B K40Q insert using primers with restriction sites (underlined):

BglII *α*Tub84B-F; 5’-TAC<u>AGATCT</u>ATGCGTGAATGTATCTCTATCCATG-3’,

NotI *α*Tub84B-R; 5’-<u>GCGGCCGC</u>TTCTGCTATACGTGTCTTTGTGGATAA-3’. Then the amplified fragment was subcloned into BglII/NotI sites of pUAST-GFP-attB vector (Kind gift from Cheng-Yu-Lee) and sequenced. Transgenic flies were generated using strain 24482 by PhiC31 integrase-mediated transgenesis (BestGene).

#### UASp-GFP-Lky

Lky cDNA (CG17003) is an intron-less gene. We amplified cDNA from yw genomic DNA using the following primers with restriction sites (underlined):

BamHI LkyF 5’-TC<u>GGATCC</u>Gatggtggagttcgcctttgaca-3’
AscI LkyR 5’-TA<u>GGCGCGCC</u>ttagaatctccggcccccggaaacc-3’

PCR product was then digested with BamHI and AscI and ligated to a modified pUASP- attB-GFP vector (kind gift from Michael Buszczak) using BamHI and AscI sites located 3’ end of GFP. Transgenic flies were generated using strain attP2 by PhiC31 integrase-mediated transgenesis (BestGene).

#### UASp-mEOS

mEOS3.2 fragment was amplified using primers with restriction sites (underlined);

NotI-EOS-F; 5’-TC<u>GCGGCCGC</u>CCCCTTCACCATGagtgcgattaagccagac-3’,

AscI-EOS-R; 5’-ACT<u>GGCGCGCC</u>tcgtctggcattgtcaggcaa-3’ from mEos3.2-C1 vector (kind gift from Michael Buszczak) and subcloned into NotI/AscI sites of a modified pUASP- attB-GFP vector (kind gift from Michael Buszczak). Transgenic flies were generated using strain 24749 by PhiC31 integrase-mediated transgenesis (BestGene).

### Immunofluorescent staining

Ovaries were dissected into 1× phosphate-buffered saline (PBS) and fixed in 4% formaldehyde in PBS for 30-60 minutes, then washed three times in PBS + 0.3% TritonX-100 (PBST) for one hour, then incubated in primary antibodies in 3% bovine serum albumin (BSA) in PBST at 4°C overnight. Samples were then washed three times in PBST for one hour (three 20 minute washes), then incubated in secondary antibodies in 3% BSA in PBST for 2-4 hours at room temperature, or at 4°C overnight. Samples were then washed three times in PBST for one hour (three 20 minute washes), then mounted using VECTASHIELD with 4,6-diamidino-2-phenylindole (DAPI) (Vector Lab, H-1200).

For anti-Lky immunofluorescence, ovaries were dissected in phosphate-buffered saline (PBS), then fixed for 10 minutes in 90% methanol, 3.7% paraformaldehyde, pre-chilled at −80°C. After fixation, the steps for immunofluorescent staining were followed as described above.

The primary antibodies used were as follows: anti Hts (1B1), anti-Orb (1:20), anti rat-Vasa (1:20), and anti-Arm (1:20) were obtained from the Developmental Studies Hybridoma Bank (DSHB), anti-acetylated microtubule antibody (6-11, b-1) was obtained from Sigma (1:200), anti-Lky (1:1000) (gift from YM. Yamashita), anti-TJ[68] (1:5000) (gift from D. Godt, University of Toronto, Toronto, ON, Canada), and anti-rabbit Vasa (1:200) (Santa Cruz Biotechnology).

AlexaFluor-conjugated secondary antibodies were used at a dilution of 1:400. Images were taken using a Zeiss LSM800 confocal microscope with a 63 ×oil immersion objective (NA=1.4) and processed using Zen software or Fiji.

### Colcemid treatment

Ovaries were dissected into Schneider’s Drosophila media and colcemid (Sigma) was added to the media to a final concentration of either 10 or 100 μM. The ovaries were incubated in colcemid for 30 minutes at room temperature, then washed twice with PBS before fixation in 4% paraformaldehyde for 30 minutes. Ovaries were then stained by immunofluorescence as described above.

### Microtubule staining

To better preserve microtubules during fixation, ovaries were dissected in 60 mM PIPES, 25 mM HEPES, 10 mM EGTA, 4 mM MgSO4, pH 6.8 (PEM buffer) and fixed in 4% formaldehyde in PEM for 12 minutes. Samples were washed in phosphate-buffered saline (PBS) containing 0.3% Triton X-100 (PBST) overnight at 4°C, followed by incubation with primary antibodies at 4°C overnight. Samples were washed for 60 minutes (three 20-minute washes) in PBST at 25°C, incubated with Alexa Fluor-conjugated secondary antibodies in PBST containing 3% bovine serum albumin (BSA) at 25°C for 2 hours, washed as above, and mounted in VECTASHIELD with DAPI (Vector Labs, H-1200).

### Fluorescent in situ hybridization

Fluorescent in situ hybridization was performed as described previously [69]. Briefly, ovaries were dissected in 1X PBS and then fixed in 4% formaldehyde/PBS for 45 minutes. After fixing, they were rinsed 2 times with 1X PBS, then resuspended in 70% EtOH, and left overnight at 4°C. The next day, testes were washed briefly in wash buffer (2X SSC and 10% deionized formamide), then incubated overnight at 37°C in the dark with 50 nM of Quasar 570 labeled Stellaris probe against entire nos 3’UTR sequence (LGC Biosearch Technologies) in the Hybridization Buffer containing 2X SSC, 10% dextran sulfate (MilliporeSigma), 1 μg/μl of yeast tRNA (MilliporeSigma), 2 mM vanadyl ribonucleoside complex (NEB), 0.02% RNAse-free BSA (ThermoFisher), and 10% deionized formamide. On the third day, testes were washed 2 times for 30 minutes each at 37°C in the dark in the prewarmed wash buffer (2X SSC, 10% formamide) and then resuspended in a drop of VECTASHIELD with DAPI.

### Photobleaching

Ovaries were dissected from flies three days after eclosion in 1 mL of prewarmed (25°C) Schneider’s Drosophila medium supplemented with 10% fetal bovine serum and glutamine– penicillin–streptomycin. Hoechst 33342 (2 μg/ml) was added as necessary prior to imaging. Dissected ovaries were placed onto ‘Gold Seal™ Rite-On™ Micro Slides two etched rings’ with a drop of media, then covered with coverslips. An inverted Zeiss LSM800 confocal microscope with a 63 ×oil immersion objective (NA=1.4) was used for imaging. Photobleaching of mEOS was accomplished using a Zeiss LSM800 confocal laser scanning microscope with a 63X/1.4 NA oil objective. Zen software was used for the programming of each experiment. Laser powers and iteration were optimized to achieve an approximately 70%-100% of bleaching, a 488 nm laser (100% laser power with 10 iterations) was used. The distribution change of fluorescence was monitored every 10 seconds for the indicated duration.

### Statistical analysis and graphing

No statistical methods were used to predetermine sample size. The experiment values were not randomized. The investigators were not blinded to allocation during experiments and outcome assessment. Statistical analysis and graphing were performed using GraphPad prism 7 software. Data are shown as means ± s.d. The P values from Student’s t-test or adjusted P values from Dunnett’s multiple comparisons test are provided; shown as *P<0.05, **P<0.01, ***P<0.001, ****P<0.0001; NS, non-significant (P≥0.05).

## Acknowledgements

We thank Yukiko Yamashita, Michael Buszczak for reading our manuscript and valuable discussions. Yukiko Yamashita, Michael Buszczak, Jill Wildonger, Dena Johnson-Schlitz, Ruth Lehmann and the Bloomington *Drosophila* Stock Center and the Developmental Studies Hybridoma Bank for reagents. Christopher Bonin for manuscript editing. This research is supported by 1R35GM128678-01 from National Institute for General Medical Sciences and start-up funds from UConn Health (to M.I.).

## Author Contributions

M.I, T.S and M.A; conception and design, acquisition of data, analysis and interpretation of data, M.B.B; Generation of Lky knock down lines. All authors wrote and edited the manuscript.

---

**Movie S1-4**

Time lapse images for Figures 2H, 2J S2A and 3F respectively. Images were taken every 10 seconds.

