## Supplemental Figures for "Acetylated microtubules are required for maintenance of the barrier between two adjacent tissues"

**Figure S1**

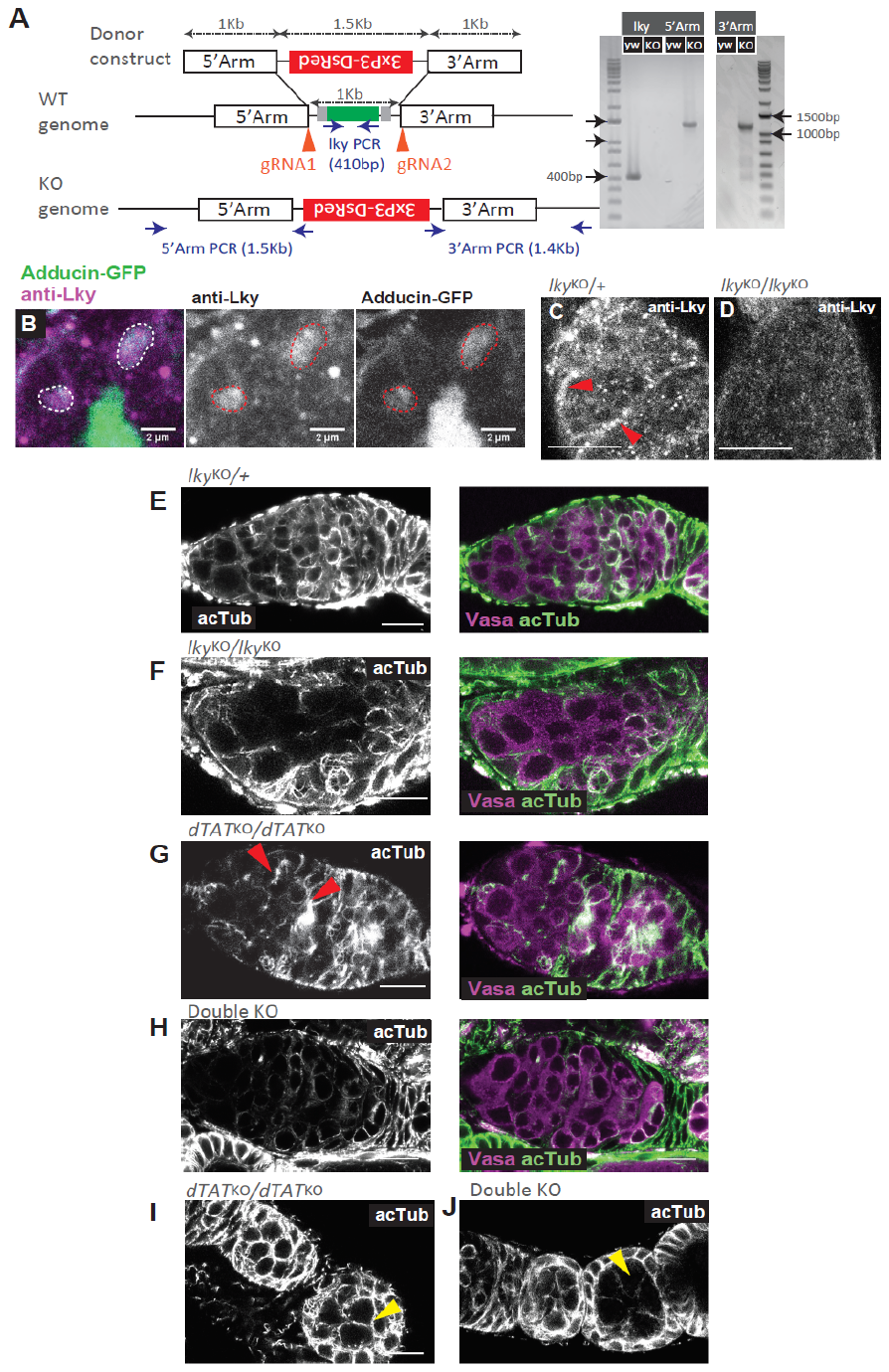

**A**) (Left) A schematic of the construct used to generate *lky*^KO^ flies. (Right) Validation of the KO genotype by PCR. **B**) IF staining with anti-Lky in flies expressing Adducin-GFP, a fusome marker. **C-D)** Anti-Lky staining in control versus KO. **E-H)** IF staining for acTub and Vasa of germaria of the indicated genotypes. “double KO” = *lky^KO^/lky^KO^;; dTAT^KO^/dTAT^KO^.* **I-J**) A comparison of acTub in early germline cysts of *dTAT^KO^* versus *Double KO*. Scale Bars: 10 µm.

**Figure S2**

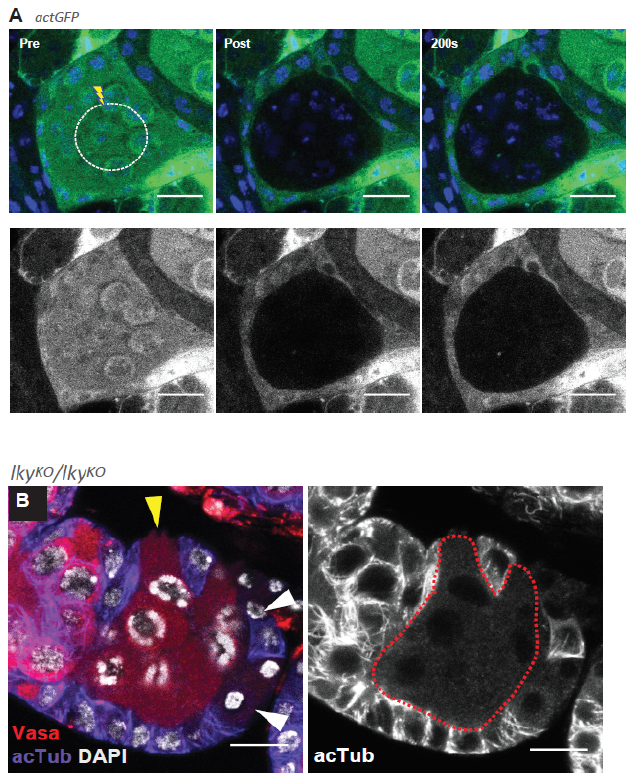

**A**) As a control for **2H**, **J** GFP was expressed ubiquitously (under the act5c promoter) in wild type flies and photobleached in the germline cyst. No recovery was observed in the germline cyst, indicating that germline and FCs do not share contents under normal conditions. **B**) A representative image of a *lky*^KO^ egg chamber stained for Vasa and acTub, showing a germline cyst extending into the somatic FC layer (yellow arrowhead) and Vasa leakage into FCs (white arrowheads). Red broken line in right panel encircles germline cyst. Scale Bars: 10 µm.

**Figure S3**

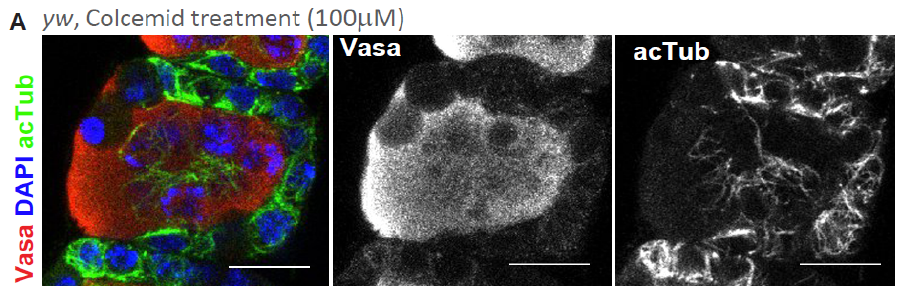

**A**) Representative image of a stage2 egg chamber after colcemid treatment, stained for Vasa and acTub, showing an acTub negative germline cyst extending into the somatic follicle cell layer. Scale Bars: 10 µm.

**Figure S5**

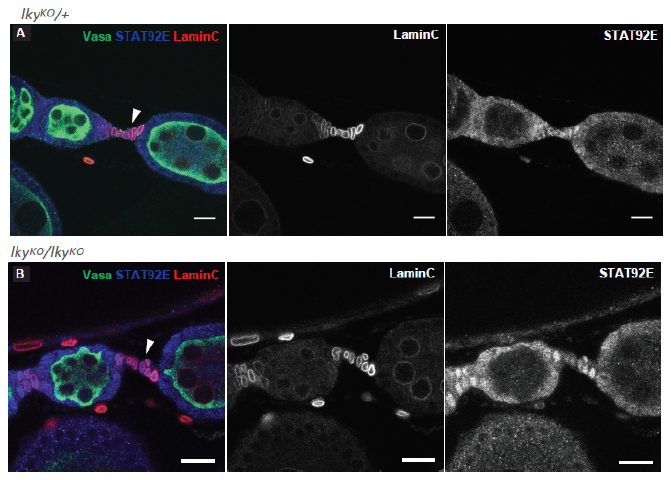

**A-B)** IF staining for Vasa, Stat92e, and LaminC, showing stalk cells marked by LaminC and Stat92e expression in control **(A)** and *lky^KO^* **(B)**. . Scale Bars; 10µm.
